# The *Arabidopsis* immune receptor EFR increases resistance to the bacterial pathogens *Xanthomonas* and *Xylella* in transgenic sweet orange

**DOI:** 10.1101/2021.01.22.427732

**Authors:** Letícia Kuster Mitre, Natália Sousa Teixeira-Silva, Katarzyna Rybak, Diogo Maciel Magalhães, Reinaldo Rodrigues de Souza-Neto, Silke Robatzek, Cyril Zipfel, Alessandra Alves de Souza

**Author notes:** Corresponding author: *Alessandra Alves de Souza,. These authors contributed equally to this work (Letícia Kuster Mitre, Natália Sousa Teixeira-Silva).

## Abstract

Plants employ cell surface receptors to recognize pathogen (or microbe)-associated molecular patterns (PAMPs/MAMPs), which are crucial for immune system activation. The well-studied *Arabidopsis thaliana* ELONGATION FACTOR-TU RECEPTOR (EFR) recognizes the conserved bacterial PAMP EF-Tu, and the derived peptides elf18 and elf26. The interfamily transfer of EFR has been shown to increase disease resistance in several crops, such as tomato, rice, wheat, and potato. Here, we generated sweet orange (*Citrus sinensis*) transgenic lines expressing *EFR* to test if it would confer broad-spectrum resistance against two important citrus bacterial diseases: citrus canker and citrus variegated chlorosis (CVC). Independent EFR transgenic lines gained responsiveness to elf18 and elf26 peptides from *Xanthomonas citri* and *Xylella fastidiosa*, as measured by reactive oxygen species (ROS) production, mitogen-activated protein kinase (MAPK) activation and defense gene expression. Consistently, infection assays showed that *Citrus-EFR* transgenic plants were more resistant to citrus canker and CVC. Our results show that the EFR immune receptor can improve plant immunity in a perennial crop against bacterial pathogens, opening perspectives to engineer durable broad-spectrum disease resistance under field conditions.

## Introduction

Plants evolved a sophisticated and highly efficient immune system as a protective mechanism against potential pathogens from the surrounding environment. Receptor kinases (RKs) and receptor-like proteins (RLPs) anchored to the cell surface function as pattern recognition receptors (PRRs) (Macho and Zipfel, 2014). These receptors sense pathogen-/microbe-/damage-associated molecular patterns (PAMPS/MAMPs/DAMPs), which are the initial event to activate pattern-triggered immunity (PTI) (Wan *et al.*, 2019). RKs are composed of an extracellular ligand-binding domain, a single-pass transmembrane domain, and a cytoplasmic kinase domain, whereas RLPs share the same basic structure, but lack a kinase domain (Boutrot and Zipfel, 2017).

PRRs recognize a large variety of PAMPs (proteins, carbohydrates, or lipids) that are generally conserved molecules essential for microbe survival (Boutrot and Zipfel, 2017). After elicitor recognition, defense responses are activated to prevent pathogen establishment and disease progression, including the production of reactive oxygen species (ROS), callose deposition, activation of Ca^2+^-dependent protein kinases and mitogen-activated protein kinases (MAPKs), and induction of defense genes (Zipfel and Oldroyd, 2017).

Several members of the leucine-rich repeat (LRR)-RK subfamily XII are PRRs recognizing bacterial proteinaceous PAMPs. Among them, the *Arabidopsis thaliana* (hereafter *Arabidopsis*) FLAGELLIN SENSING 2 (FLS2) and EF-TU RECEPTOR (EFR) are the receptors for bacterial flagellin (or the derived epitope flg22) and EF-Tu (or the derived epitopes elf18 and elf26) (Gómez-Gómez and Boller, 2000; Zipfel *et al.*, 2006). While FLS2 is present in all higher plants, some plant species have developed additional receptors for flagellin. The FLS3 receptor present in *Solanaceae* plants recognizes the flagellin epitope flgII-28 (Hind *et al.*, 2016), while rice perceives the *Acidovorax avenae* flagellin via a yet-unknown receptor (Katsuragi *et al.*, 2015). Different from FLS2, EFR is restricted to the *Brassicaceae* family (Zipfel *et al.*, 2006). However, another fragment in the middle region of EF-Tu (termed Efa50) can be recognized via a yet-unknown receptor in some rice varieties (Furukawa *et al.*, 2014).

Genetic transfer of PRRs has been shown to be effective in conferring broad-spectrum resistance in several crop species, demonstrating that downstream signaling components are widely conserved even among phylogenetically distant species (Boutrot and Zipfel, 2017; Rodriguez-Moreno *et al.*, 2017). Particularly, EFR from *Arabidopsis* was reported to confer resistance in *Nicotiana benthamiana* and tomato against phytopathogenic bacteria belonging to different genera (Kunwar *et al.*, 2018; Lacombe *et al.*, 2010). Rice plants expressing *EFR* also showed increased resistance against two pathogenic bacteria, *Xanthomonas oryzae* pv. *oryzae* and *A. avenae* subsp. *avenae* (Holton *et al.*, 2015; Lu *et al.*, 2015; Schwessinger *et al.*, 2015). Transgenic *EFR*-expressing wheat demonstrated enhanced resistance to *Pseudomonas syringae* pv. *oryzae* (Schoonbeek *et al.*, 2015), and more recently, potato and *Medicago truncatula* expressing EFR showed greater resistance to *Ralstonia solanacearum* (Boschi *et al.*, 2017; Pfeilmeier *et al.*, 2019).

Bacterial diseases have been associated with major economic losses in commercial citrus orchards. Especially, Brazil, the biggest sweet orange (*Citrus sinensis*) producer, faces serious problems to manage citrus bacterial pathogens (Neves *et al.*, 2020). Among them, citrus canker and citrus variegated chlorosis (CVC), caused by *X. citri* subsp. *citri* and *Xylella fastidiosa* subsp. *pauca*, respectively, are important bacterial diseases and are therefore extensively studied due to their impact on citrus agribusiness (Caserta *et al.*, 2020; Coletta-Filho *et al.*, 2020; Martins *et al.*, 2020). All sweet orange (*Citrus sinensis*) commercial varieties are susceptible to both diseases and, despite many efforts, no natural resistance has been found in *C. sinensis* so far. The most effective approach for citrus breeding is based on gene introgression from close relatives; however, obtaining varieties carrying durable resistance combined with desirable agronomic traits can be challenging and time-demanding (Machado *et al.*, 2011). Thus, biotechnological approaches are a powerful strategy to support programs in increasing resistance to biotic stresses (Caserta *et al.*, 2020). Since EF-Tu is present in the biofilm of both bacteria (Silva *et al.*, 2011; Zimaro *et al.*, 2013) and the outer membrane vesicles (OMVs) released by *X. fastidiosa* (Nascimento *et al.*, 2016), we hypothesized that *EFR* gene transfer as an attempt to conferring broad-spectrum resistance in citrus plants shows great potential.

Here, we generated transgenic sweet orange expressing *EFR*. The transgenic lines were able to sense elf peptides from *X. citri* and *X. fastidiosa*, as measured by ROS production, MAPK activation and defense marker gene expression. This activation and signaling of the citrus immune system culminated in reduced symptom development and increased resistance to citrus canker and CVC. This strategy provides commercial sweet orange varieties harboring important agronomic traits with PRR-based resistance against bacterial diseases to better support future needs of citrus breeding programs.

## Material and Methods

### Vectors and plant genetic transformation

The binary vector containing the *EFR* gene from *Arabidopsis* was chemically synthesized by the DNA Cloning Service e.K. company (www.dna-cloning.com/). The transgene is under the control of the Figwort Mosaic Virus (FMV) promoter and the *Agrobacterium* nopaline synthase (NOS) terminator. The vector carries kanamycin resistance on the T-DNA, streptomycin/spectinomycin-resistance for bacterial selection, and *gus* reporter gene (Fig. S1).

*Agrobacterium*-mediated transformation was used to produce citrus transgenic lines as previously described by Caserta *et al.* (2014). Seeds were sampled from mature Valencia sweet orange fruits and cultured in MS/2 solid medium for four weeks in the dark at 27 °C. Seedlings of about 15 cm in length were transferred to a 16-h photoperiod for 15 days and later used as explant source for genetic transformation. Epicotyl segments (0.8 - 1.0 cm) were excised and kept in liquid MS medium supplemented with indole acetic acid (100 mg L^−1^) before incubation in *Agrobacterium tumefaciens* (EHA105 strain) suspension (10^8^ CFU mL^−1^) for 5 minutes. Dried explants were transferred to solid MS co-culture medium supplemented with sucrose (30 g L^−1^), benzylaminopurine (BAP) (10 mg L^−1^), and myo-inositol (100 mg L^−1^) for three days (24 ⁰C, in the dark). After the co-culture period, the explants were transferred to solid MS selection medium containing kanamycin (100 mg L^−1^) and cefotaxime (250 mg L^−1^) for four weeks (28 ⁰C, in the dark) and then retransferred to 16-h photoperiod.

Kanamycin-resistant shoots were excised from the explants and incubated in 2mM X-Gluc solution (37 °C, overnight) for β-glucuronidase (*gus*) assay. Genomic DNA was extracted from leaves using the CTAB method (Doyle and Doyle, 1990) and PCR was performed with the annealing of the forward primers to the FMV promoter (FMV_F) and the reverse primer within the *EFR* sequence (*EFR*_R) (Table S1). The well-developed and PCR positive shoots grown *in vitro* were directly grafted on Rangpur lime (*Citrus limonia*) rootstocks and kept under greenhouse conditions. Source plants were used to produce clones for subsequent evaluations. Transformation efficiency was calculated as the percentage of *gus*-positive shoots in the total of explants exposed to *Agrobacterium* culture.

*Nicotiana tabacum* was used as an experimental model to assess the role EFR plays in recognizing citrus bacterial PAMPs and its response to *X. fastidiosa* infection. The binary vector pEarlyGate103 containing the open reading frame of *Arabidopsis EFR* was used (Lacombe *et al.*, 2010; Zipfel *et al.*, 2006). Transgenic tobacco was produced by *Agrobacterium*-mediated transformation with the strain GV3101, as previously described (Gómez *et al.*, 2020). Vigorous shoots grew in selective media (MS/2 supplemented with 2 μg mL^−1^ phosphinothricin) were transferred to pots containing 2:1 substrate/vermiculite and kept in an acclimatization room at 28 °C for 21 days. The presence of the expression cassette *35S∷EFR-GFP-His* was confirmed by PCR using genomic DNA as template and the primers 35S_F and *EFR*_R (Table S1).

### ROS production assay

The peptides elf18_Ec_ (ac-SKEKFERTKPHVNVGTIG), elf18_Xcc_ (ac-AKAKFERTKPHVNVGTIG), and elf26_Xf_ (ac-AQDKFKRTKLHVNVGTIGHVDHGKTT) from *Escherichia coli*, *X. citri* and *X. fastidiosa*, respectively, were synthesized by Aminotech Research and Development. Leaf discs (0.5 cm) from young tender leaves were displaced on autoclaved water overnight in a 96-well plate at room temperature and then challenged by 100 μL of elicitation solution (17 mM luminol, 1 μM horseradish peroxidase and 100 nM elf18_Ec_, elf18_Xcc_ or elf18Xf). Luminescence was immediately measured over 40 minutes using the Varioskan Flash Multiplate Reader (Thermo Scientific). Assays were performed in triplicates and statistical significance of the means calculated according to Tukey’s test (* *p* < 0.05).

For ROS production in response to *X. fastidiosa* bacteria, petioles of six- to seven-week-old *Arabidopsis* Col-0 and *efr-1* mutant plants (Zipfel *et al.*, 2006) were sampled using a scalpel and left overnight in sterile water. The following day the water was replaced with a solution containing 17 μg mL^−1^ (w/v) luminol (Sigma), 10 μg mL^−1^ horseradish peroxidase (Sigma) and living *X. fastidiosa* subsp. *fastidosa* Temecula-1(Ionescu *et al.*, 2014) cells (OD_600_ 0.125). Luminescence was captured using a TECAN plate reader Infinite® 200 PRO.

### Gene expression analysis by RT-qPCR

Total RNA was isolated from leaf tissue of transgenic and non-transgenic plants using the RNeasy Mini Kit (Qiagen) following the manufacturer’s instructions and treated with RNase free-DNase (Promega). The RNA samples were PCR-tested for genomic DNA cross-contamination. RNA quality and concentration were assessed by gel electrophoresis and spectrophotometry (NanoDrop 8000 - Thermo Scientific). cDNA was synthesized from 1 μg total RNA using High-Capacity cDNA Reverse Transcription Kit (Applied Biosystems) using Oligo(dT)15. The relative expression values were analyzed using the SYBR Prime Script RT-PCR kit (Thermo Scientific) in ABI PRISM 7500 Fast (Applied Biosystems) and were determined by the ΔΔCt method (Livak and Schmittgen, 2001). Expression values were normalized by the endogen *cyclophilin* for *C. sinensis* and by the gene *ARPC3* (Actin-related protein C3) for tobacco. The primers used for RT-qPCR are listed in Table S1. Assays were performed in triplicates and statistical significance of the means calculated according to Tukey’s test (* *p* < 0.05).

### MAP kinase assay

Leaf discs (0.5 cm) from young tender leaves were displaced on autoclaved water overnight in a 96-well plate at room temperature and were treated with flg22_Pst_ from *Pseudomonas syringae*, elf18_Ec_, elf18_Xcc_, elf26_Xf,_ or water (mock treatment) for 0, 30, and 45 minutes and immediately frozen in liquid nitrogen. The samples were ground into powder before the addition of extraction buffer [50 mM Tris-HCl pH 7.5, 100 mM NaCl, 15 mM EGTA, 10 mM MgCl_2_, 1 mM NaF, 1 mM Na_2_MoO_4_.2H_2_O, 0.5 mM NaVO_3_, 30 mM β-glycerophosphate, 0.1 % IGEPAL CA 630, 100 nM calyculin A (CST), 0.5 mM PMSF, 1 % protease inhibitor cocktail (Sigma, P9599) and 5 % glycerol]. The extracts were centrifuged at 16,000 x g and 5x SDS loading buffer added. Protein concentrations were measured by the Bradford assay (Protein Assay Dye Reagent - Bio-Rad) and 30 μg of total protein was separated by 12 % SDS-PAGE and blotted onto PVDF membrane (Bio-Rad). The membranes were blocked in 5 % (w/v) BSA (Sigma) in TBS-Tween (0.1 %) for 1 hour. The activated MAP kinases were detected using anti-p42/44 MAPK primary antibody (1:2500, Cell Signaling Technology, 4370) overnight at 4 °C, followed by anti-rabbit HRP-conjugated secondary antibody (Sigma). Three independent experiments were performed with similar results.

### Seedling growth assay

*Arabidopsis* Col-0, *efr-1* and *bak1-5* mutant seeds were surface-sterilized, sown on MS media supplemented with sucrose, stratified for 2 days at 4 °C in the dark and put in the light. Four days-old seedlings were transferred into liquid MS with or without *Xff*OMVs (see below) and incubated for ten further days. Fresh weight of seven replicates per treatment and genotype was measured using a precision scale and calculated relative to untreated control.

OMVs from *Xylella fastidiosa* subsp. *fastidiosa* Temecula-1 were isolated as described in Ionescu *et al.* (2014) with some modifications. Briefly, *X. fastidiosa* subsp. *fastidiosa* Temecula-1 was cultured in 100 mL of PD2 medium for seven days. Cells were removed by centrifugation at 10,000 × g for 15 min at 4 °C. The supernatant was filtered through 0.22 μm filter and centrifuged at 38,000 × g for 1 h at 4 °C. The supernatant was removed carefully and subjected to centrifugation at 150,000 × g for 4 h at 4 °C. The final pellet containing OMVs was resuspended in 1 mM EDTA (pH 8.0) and stored frozen at −80 °C until used.

### *Xanthomonas citri* infection assay

*X. citri* subsp. *citri* strain 306 (Da Silva *et al.*, 2002) expressing GFP (Rigano *et al.*, 2007) was grown overnight in liquid NBY medium supplemented with ampicillin (100 mg L^−1^) and gentamicin (5 mg L^−1^). The bacterial suspension (10^4^ CFU mL^−1^) was prepared in 1x phosphate-buffered saline (PBS) and infiltrated in three regions of three transgenic and non-transgenic fully expanded detached leaves. Citrus canker symptoms were evaluated 7 and 14 days after inoculation (dai). Leaf discs adjacent to the infiltration site were excised to assess the bacterial population from three independent leaves. This assay was performed in triplicates and the statistical significance of the means calculated according to the Student’s *t*-test (* *p* < 0.05).

### *Xylella fastidiosa* infection assay

*X. fastidiosa* subsp. *pauca* strain 9a5c (Simpson *et al.*, 2000) was cultivated in solid periwinkle wilt medium (PWG) (Davis *et al.*, 1981) for 7 days at 28 °C. The bacterial suspension (10^8^ CFU mL^−1^) was prepared in 1x PBS buffer for petiole inoculation on the first leaf of ten transgenic and wild-type (WT) plants, for both transgenic tobacco and citrus. One month after inoculation, genomic DNA was extracted from petioles of the first leaf above the inoculation point (aip), as previously described, and bacterial detection was performed by PCR using RST31/33 primers (Minsavage *et al.*, 1994). Only the *Xylella*-positive and mock-inoculated individuals were maintained for further analysis.

Transgenic citrus plants were evaluated regarding *X. fastidiosa* population 18 months after inoculation by qPCR in ABI PRISM 7500 Fast (Applied Biosystems). Petioles were collected in two different parts of the plants for total DNA extraction, at 5 and 30 cm aip. The quantification was performed using TaqMan PCR Master Mix (Applied Biosystems) using primers CVC-1 and CCSM-1 and the probe TAQCVC (Table S1) and the bacterial population was calculated according to the standard curve developed for *X. fastidiosa* (Oliveira *et al.*, 2002). Infected plants were evaluated for the disease severity and scored by three inspectors using a diagrammatic scale (Caserta *et al.*, 2017).

Three independent evaluations of the infected *EFR*-expressing tobacco plants were performed by three inspectors and scored for disease incidence at 30, 40, 50, and 60 dai. The disease incidence corresponds to the proportion of symptomatic leaves by the total number of leaves multiplied by 100. The area under the disease progress curve (AUDPC) was calculated based on the trapezoidal integration model (Berger, 1988), according to the equation: *AUDPC = (yi + ys) ÷ 2 × t*, where *yi* refers to the mean of the incidence value given to symptoms at a given time point and *ys* to the mean of the incidence value given to the immediately following time point, and *t* to the interval of time of each evaluation. Statistical significance of the means was evaluated according to the Student’s *t*-test (* *p* < 0.05; ** *p* < 0.01).

*Arabidopsis* Col-0 and *efr-1* plants were infected with *X. fastidiosa* subsp. *fastidiosa* Temecula-1 as described in Pereira *et al.* (2019). Briefly, twelve six- to seven-week-old plants per genotype were inoculated by dropping 5 μL of bacterial inoculum or PBS buffer at the midrib. The petiole tissue under the drop was pricked seven to eight times using an insulin needle. After 14 days, petiole tissue below the infection point was harvested for DNA isolation and qPCR of the *HL* gene from *X. fastidiosa* subsp. *fastidiosa* Temecula-1.

## Results

### EFR is responsive to elf peptides from *X. citri* and *X. fastidiosa*

We first examined the ability of EFR to respond to *X. fastidiosa* bacterial suspension. *Arabidopsis* Col-0 plants produced a prototypic ROS burst when challenged with living *X. fastidiosa* (Fig. 1a; Fig. S2a). This ROS burst was markedly reduced in *efr-1* mutants, suggesting the perception of immunogenic elf peptides from *X. fastidiosa* EF-Tu. Since EF-Tu is present in outer membrane vesicles (OMVs) released from *X. fastidiosa* (Feitosa-Junior *et al.*, 2019), we determined the effect of OMVs on *Arabidopsis* seedling growth, which is typically inhibited by continual PAMP treatment (Zipfel *et al.*, 2006). The growth of *Arabidopsis* Col-0 seedlings was strongly repressed in the presence of *X. fastidiosa* OMVs (Fig. 1b). No growth repression was observed in *efr-1* or *bak1-5*, a mutant affected in the EFR co-receptor BRASSINOSTEROID INSENSITIVE 1-ASSOCIATED RECEPTOR KINASE 1 (BAK1) involved in PTI (Schwessinger *et al.*, 2011). We then examined whether EFR modulates the success of *X. fastidiosa* infection in *Arabidopsis* (Pereira *et al.*, 2019). Compared to wild-type Col-0 plants, *efr-1* mutants supported higher bacterial loads of *X. fastidiosa*, quantified as *HL* gene abundance (Fig. 1c). Thus, perception of EF-Tu (and potentially derived elf peptides) by EFR is sufficient to restrict *X. fastidiosa* colonization in *Arabidopsis*.

**Fig. 1.**
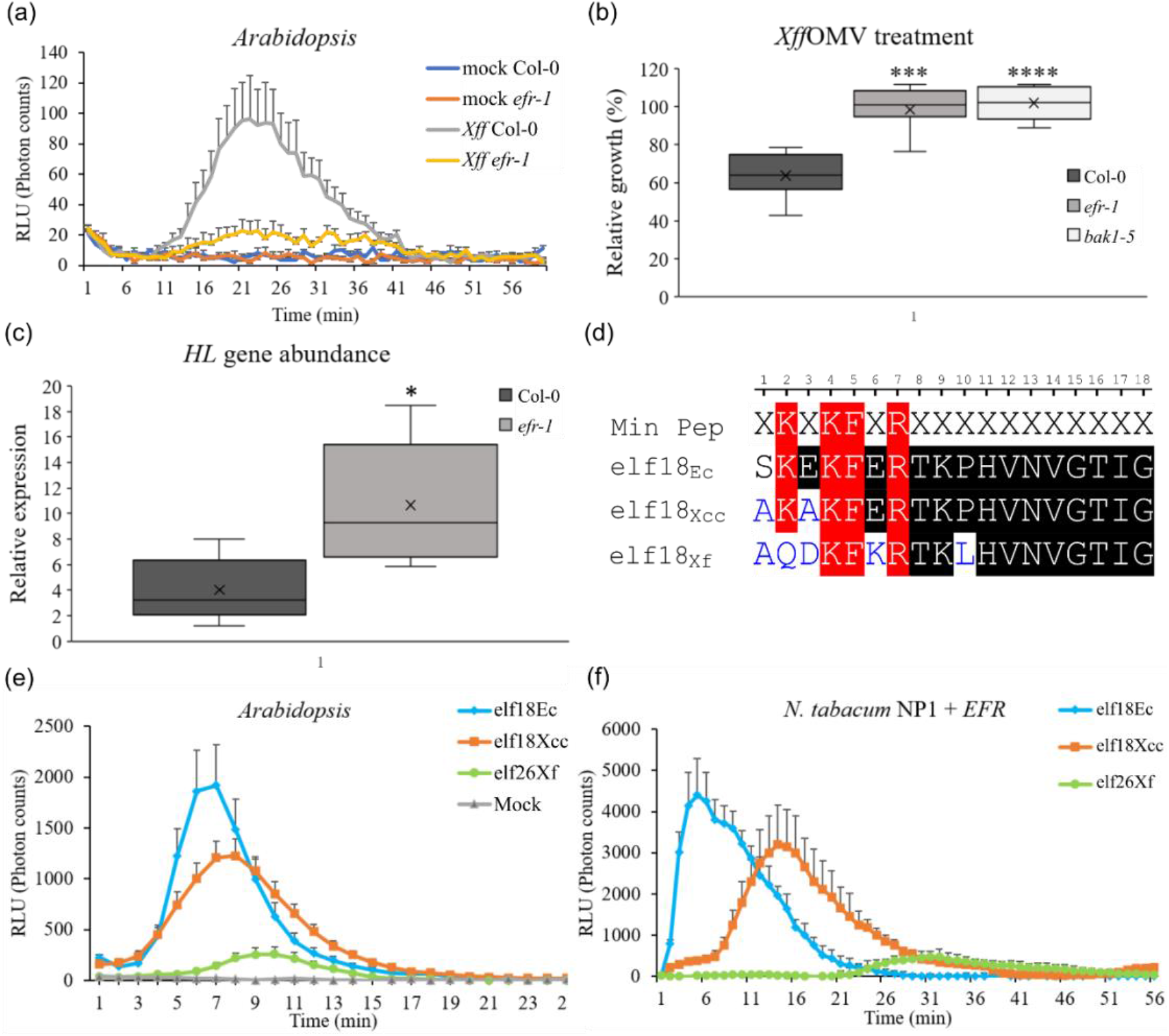
The perception of *X. fastidiosa* and elf peptides in *Arabidopsis* (Col-0 and *efr-1*) and transgenic tobacco expressing *EFR*. **(a)** *efr-1* mutant is strongly impaired in *Xff*-induced ROS burst. Average photon count, represented as RLU, over 60 min following *Xff* treatment (n=10). **(b)** Seedling growth inhibition of Col-0, *efr-1* and *bak1-5* in the presence of *Xff*OMVs (1:50 dilution). Fresh weight is represented relative to untreated control. Results are average +/− SEM (n=7) **(c)** Relative expression of *Xylella HL* in *Arabidopsis* Col-0 and *efr-1* after infection by bacterial suspension (n=5). **(d)** Alignment of the EF-Tu-derived elf18 sequences from *E. coli*, *X citri* and *X. fastidiosa* compared to the minimal peptide (in red, where X is any amino acid) required for full EFR elicitation. Amino acids in blue represent substitutions in the respective peptide sequences. **(e)** ROS production in *Arabidopsis* (Col-0) triggered by elf peptides from *E. coli* (elf18_Ec_), *X. citri* (elf18_Xcc_) and *X. fastidiosa* (elf26_Xf_) (n=6). **(f)** Representation of ROS production in transgenic tobacco expressing *EFR* after treatment with elf peptides (n=6). RLU: relative light units. Values are means ± standard error (SE). Statistical differences were calculated using a two-tailed t-test (* *p* < 0.05). The experiments were performed three times with similar results.

Prompted by these results, we next investigated whether the elf peptides derived from the citrus pathogenic bacteria *X. fastidiosa* and *X. citri* activate downstream responses upon perception by EFR. We tested ROS activation in *Arabidopsis* (Col-0) and in transgenic tobacco expressing *EFR*. We generated five transgenic tobacco lines (T1, T2, T3, T4, and T5) in which *EFR* integration and expression were confirmed (Fig. S3a, b). Since *Arabidopsis* (Col-0) naturally harbors EFR, the activation of ROS production quickly occurred after elf18_Ec_ exposure (Fig. 1d, Fig S2b). Although there are sequence differences between elf peptides (Fig. 1d), ROS production was similarly observed when *Arabidopsis* leaf discs were challenged with elf18_Xcc_ (Fig. 1e, Fig S2b). However, when treated with elf26_Xf_, ROS production was comparatively delayed and significantly lower (Fig. 1e, Fig S2b). A similar pattern was observed for the transgenic tobacco plants expressing *EFR* (Fig. 1f). Altogether, these findings reinforce that EFR can recognize peptides derived from citrus bacterial pathogens, indicating that its transfer to sweet orange might be a good strategy to confer broad recognition of these bacteria.

### Citrus plants expressing EFR are responsive to elf peptides

Nine independent transgenic lines of Valencia sweet orange (V1, V2, V3, V4, V5, V6, V7, V8, and V9) were successfully transformed with *EFR* (Table S2). Transgenic lines were selected by tissue culture via kanamycin selection and histochemical *gus* assay (Fig. 2a). Transgene integration and expression were confirmed by PCR and RT-qPCR, respectively (Fig. 2b, c). The relative transcript levels varied between lines, with V4 showing the highest expression level, while V7 showed the lowest (Fig. 2c). Phenotypic abnormalities were absent after grafting in Rangpur lime rootstocks, except for the V6 line, which showed compromised leaf morphology and vegetative development (data not shown). This line was therefore omitted from further assays.

**Fig. 2.**
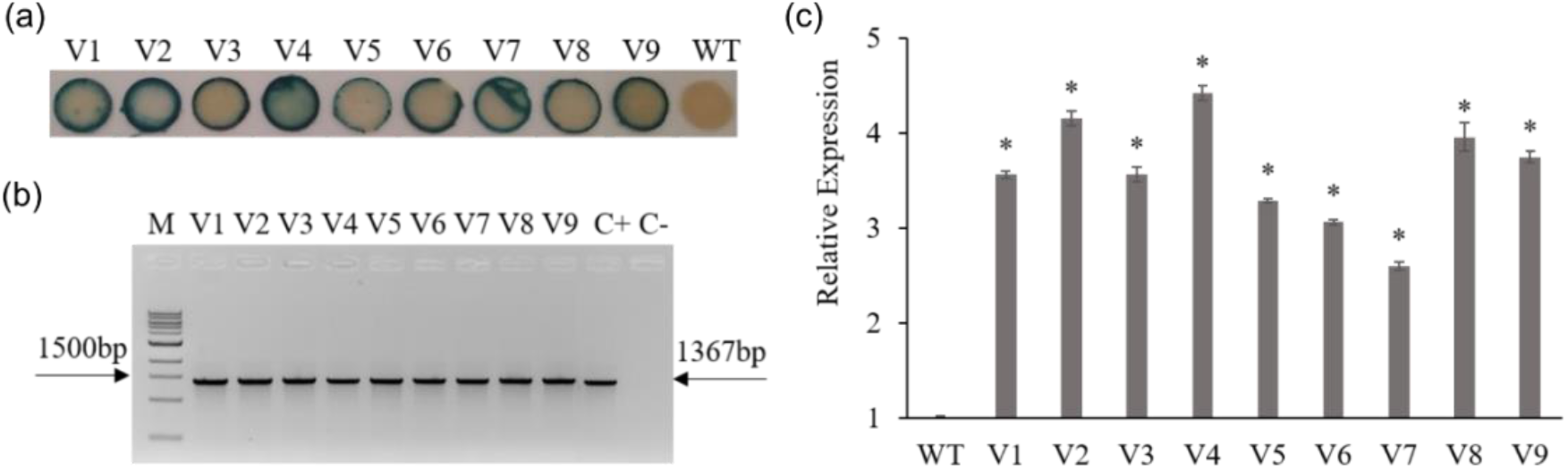
Molecular confirmation of *EFR*-expressing transgenic citrus plants. **(a)** Histochemical *gus* assay of transgenic lines and wild-type (WT). **(b)** Analysis of the PCR product (1367 bp) in 1 % agarose gel depicting the presence of the transgene in Valencia transgenic citrus lines. M: 1 kb Plus DNA Ladder (Fermentas); C+: positive control_binary vector; C-: negative control_sweet orange WT. **(c)** The relative expression level of *EFR* measured by RT-qPCR normalized by the expression of *cyclophilin*. Data are represented as the mean values ± standard error (SE) of three technical replicates. Statistical differences were calculated using the Student’s *t*-test (* *p* < 0.05). Experiments were repeated three times with similar results.

Attempting to verify the effectiveness of the sweet orange *EFR* transgenic lines in recognizing multiple elf peptides and to confirm whether EFR is functional in citrus, we challenged the transgenic plants with elf18_Ec,_ and ROS production was measured in all transgenic lines, showing variable total ROS production levels (Fig. 3a). The recognition of elf18_Ec_ and subsequent ROS production by the transgenic citrus plants indicate the functional conservation of the required intracellular signaling components in sweet orange. Similar results were obtained after treatment with the *X. citri* peptide, in which all transgenic lines were able to recognize elf18_Xcc_ and trigger ROS production (Fig. 3b). Surprisingly, though, ROS production was not easily detected in most transgenic lines after elf26_Xf_ challenging. A delayed and weaker ROS production was measured in the V4 and V5 lines (Fig. 3c). The lower elf26_Xf_ activity was evident when compared to the peaks obtained from both elf18_Ec_ and elf18_Xcc_ exposure (Fig. 3).

**Fig. 3.**
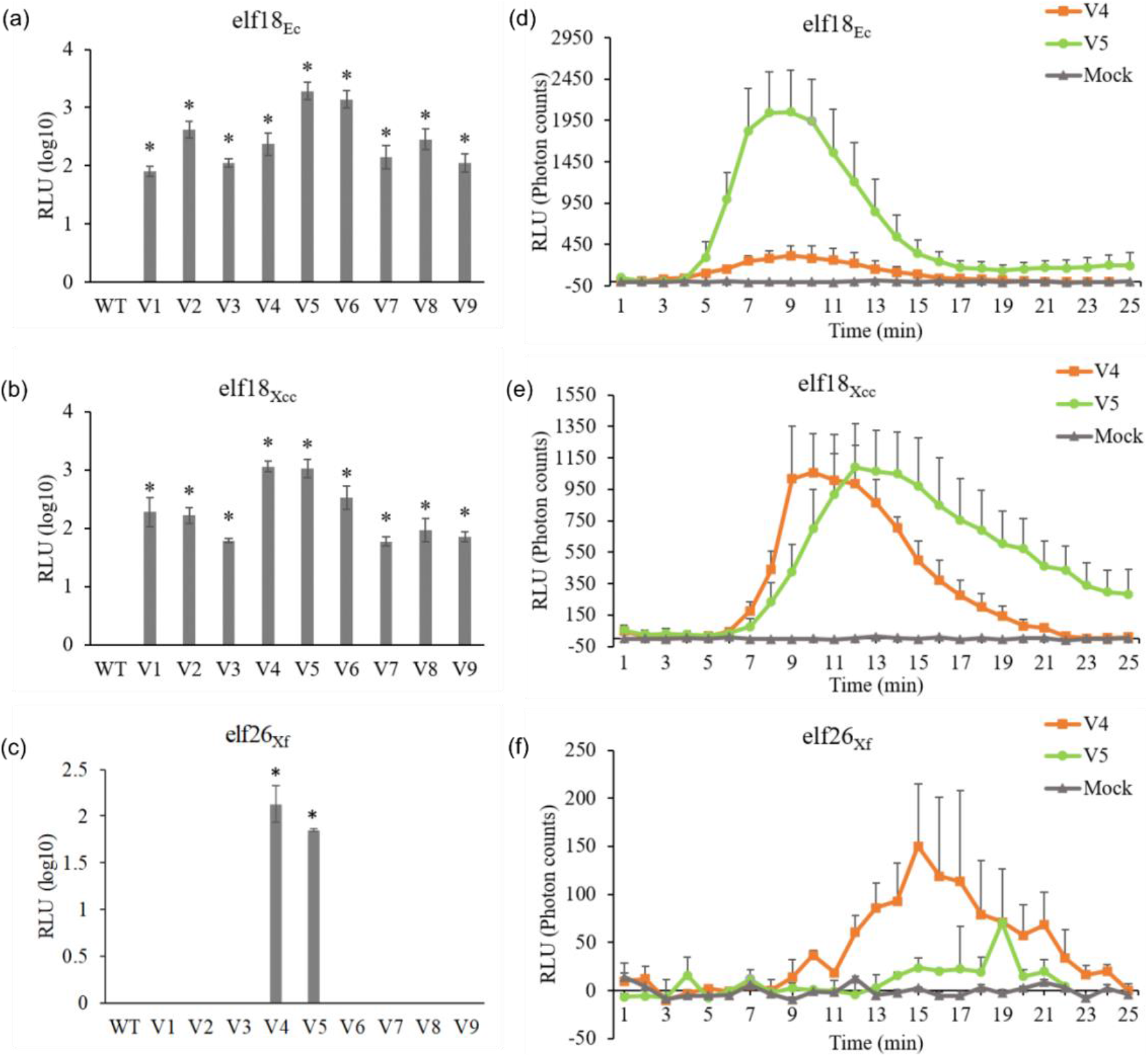
Perception of elf18_Ec_, elf18_Xcc,_ or elf26_Xf_ in *EFR* transgenic citrus plants **(a, b, c)** ROS production in response to elf peptides was verified in nine transgenic lines. **(d, e, f)** Temporal ROS production over 25 minutes in response to elf peptides in V4 and V5 transgenic lines. RLU: relative light units. Values are means ± standard error (SE) of at least six biological replicates, except to elf26_Xf_ (n=3). Statistical differences were calculated using the Student’s *t*-test (* *p* < 0.05). Experiments were repeated three times with similar results

Since V4 and V5 could sense all the three peptides herein studied and showed the best performance to activate immune responses in citrus, these lines were chosen to test ROS production over time (Fig. 3d-e). The ROS peaks occurred 9 minutes after elf18_Ec_ treatment, lasting about 17 minutes (Fig. 3d). For elf18_Xcc_ and elf26_Xf_, ROS peaks occurred after 10 and 15 minutes, respectively (Fig. 3e-f). WT citrus plants were insensitive to elf18_Ec_, elf18_Xcc_, and elf26_Xf_ (Fig. 3a-c).

### elf peptides activate MAPKs and defense-related genes in *EFR* citrus plants

To test if *EFR* expression results in the activation of downstream immune responses, MAPK activation and the expression of defense-related genes were measured. MAPK phosphorylation was detected in both V4 and V5 transgenic lines compared to the mock treatment 30 and 45 minutes after elf treatment. For both transgenic lines, MAPK showed a stronger accumulation 45 minutes after elf peptide treatment (Fig. 4). Interestingly, constitutive activation was observed in the V5 line; nevertheless, MAPK activation still increased after peptide treatment (Fig. 4).

**Fig. 4.**
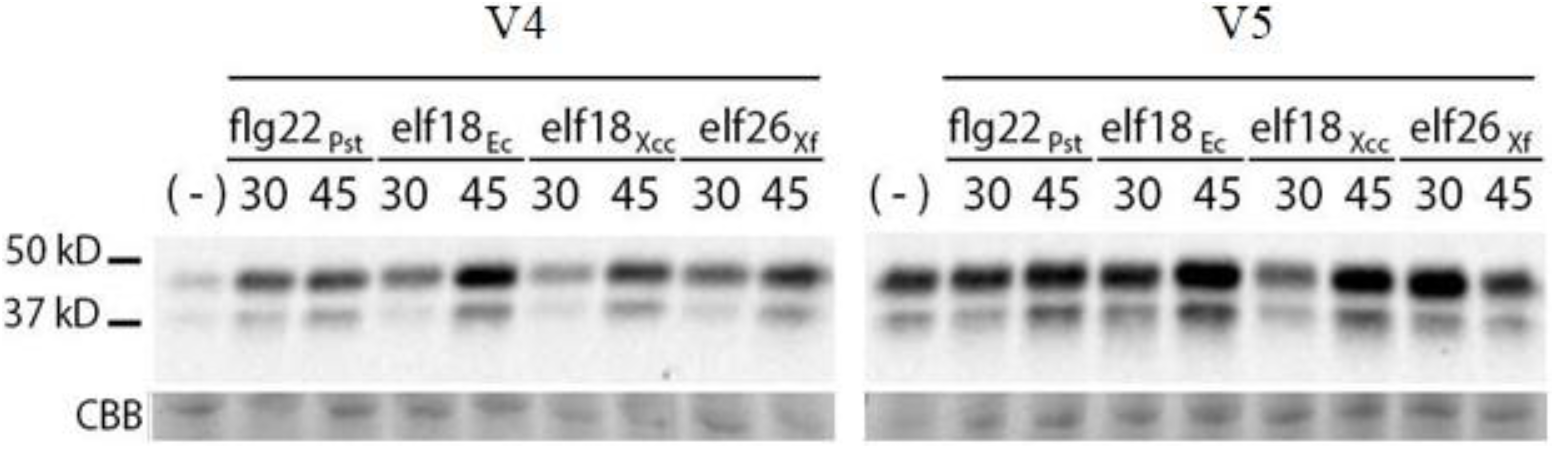
MAPK activation in transgenic sweet orange after treatment with elf peptides. The leaves were treated with elf18_Ec_, elf18_Xcc_, elf26_Xf_ or flg22_Pts_ (positive control) and water (-) and collected 30 and 45 minutes after treatment. MAPK phosphorylation was detected by western blotting using anti-phospho p44-p42-antibody. Even loading is demonstrated by Coomassie Brilliant Blue (CBB) staining. Experiments were repeated three times with similar results.

We sought to assess the expression behavior of four citrus defense marker genes (*SGT1*, *EDS1*, *WRKY23,* and *NPR2*) (Rodrigues *et al.*, 2013; Shi *et al.*, 2015). All genes were induced by elf peptides to different levels in the V4 and V5 transgenic lines compared to the mock treatment (Fig. 5). Notably, the upregulation of the genes in response to most peptides was stronger in V5 than in the V4 line. Besides, following the same pattern observed for ROS production, a stronger induction of defense genes was triggered after elf18_Ec_ and elf18_Xcc_ in comparison to elf26_Xf_ treatment (Fig. 5).

**Fig. 5.**
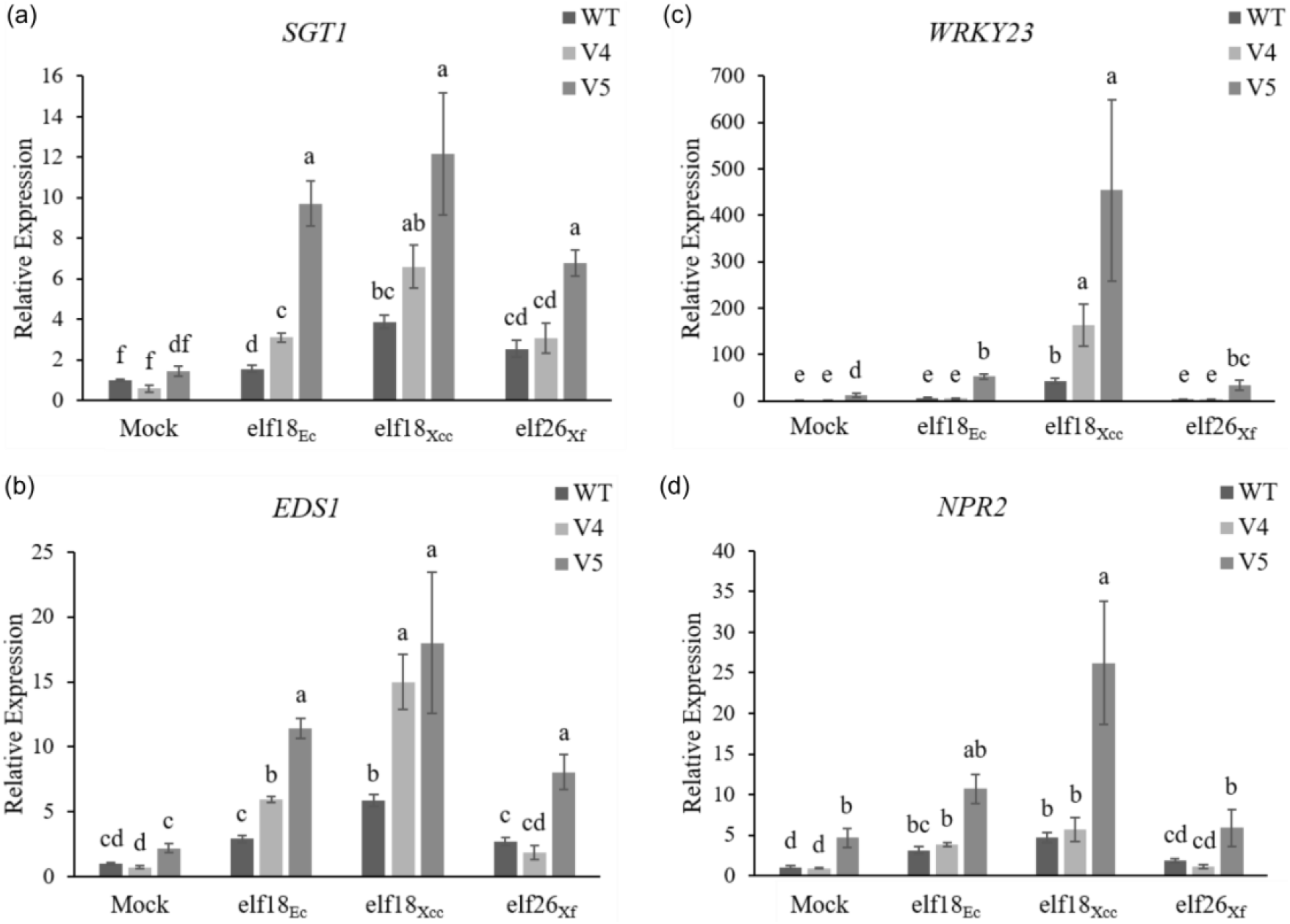
Expression pattern of defense-related genes in citrus-*EFR* transgenic lines V4 and V5. Four genes (*SGT1*, *WRKY23*, *EDS1,* and *NPR2*) were evaluated 3 hours after treatment with elf18_Ec_, elf18_Xcc_ or elf26_Xf,_ and water (mock). Relative gene expression levels were measured by RT-qPCR and normalized to the expression of *CYCLOPHILIN*. Fold change is relative to the mock treatment. Data are expressed as the mean values ± standard error (SE) of three biological replicates. Different letters on the top of the bars indicate significant statistical differences among the mean values calculated with one-way ANOVA followed by Tukey’s test (*p* < 0.05).

### Transgenic sweet orange expressing *EFR* shows enhanced resistance to citrus canker

Since the presence of EFR enabled sweet orange to recognize elf peptides derived from bacteria infecting citrus and triggered immune signaling outputs, we evaluated whether the transgenic plants show enhanced resistance to citrus canker. Detached leaves were infiltrated with *X. citri* bacterial suspension and disease progression was assessed 7 and 14 days after inoculation (dai). Transgenic lines and WT plants did not show any canker symptom at 7 dai and bacterial growth was not significantly different among the treatments (Fig. 6b). Typical canker lesions developed in all inoculated leaves 14 dai (Fig. 6). Both transgenic lines showed reduced symptom severity when compared to the WT (Fig. 6a). Notably, the V5 line produced mild hyperplasic and water-soaked lesions, and petiole abscission, an advanced stage mark of canker disease, was never observed (Fig.6a). Although reduced symptom development could be observed in V4, the bacterial population was not significantly different from the WT control. However, the pathogen growth in the V5 line was consistently lower, showing a reduction in the order of 3 log units (Fig. 6b). In this transgenic line, bacterial spreading and growth seem to be somehow restrained, thus corroborating symptomatology results (Fig. 6b).

**Fig. 6.**
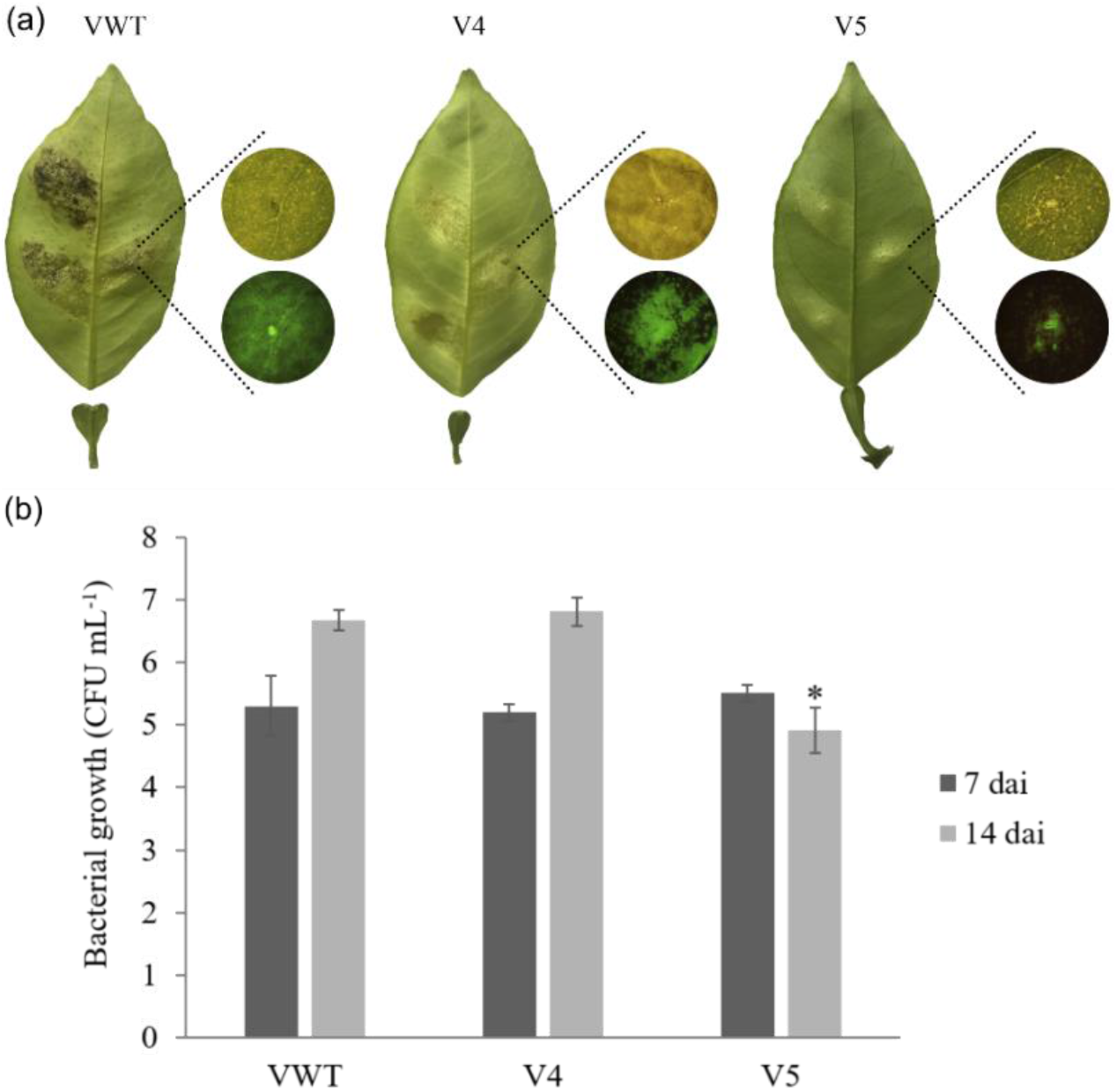
Citrus canker disease assay in transgenic sweet orange detached leaves from V4 and V5 lines, and the wild-type (WT). **(a)** Citrus canker symptomatology 14 days after *X. citri*-GFP infiltration. Circles represent the details of the area of infection in bright-field (upper circle) and the bacterial accumulation under GFP-fluorescence (lower circle). **(b)** Bacterial growth was evaluated in leaf discs adjacent to the inoculation point at 7 and 14 dai. Data are expressed in log_10_ and represent the means ± standard error of three independent lesions. Statistical differences compared to WT were determined by the Student’s *t*-test (* *p* < 0.05).

### *EFR* expression prevents *X. fastidiosa* migration and decreases CVC symptoms

To further investigate whether the presence of *EFR* can trigger immune responses upon *X. fastidiosa* infection, we first challenged transgenic tobacco expressing *EFR*. Most transgenic lines showed similar behaviors along with the disease progression, with a consistent reduction in symptoms development (Fig. S3c). To better assess the influence of the receptor in improving tolerance to *X. fastidiosa*, we calculated the area under the disease progress curve (AUDPC) based on four time-points evaluated (Fig. S3d). The transgenic trait conferred a significant reduction in the progression of *X. fastidiosa* symptoms, showing that EFR enables pathogen recognition and weakens its development within the xylem vessels (Fig. S3).

*EFR-*expressing citrus plants were also challenged with the pathogen to assess the incidence and severity of CVC. One month after inoculation, the presence of *X. fastidiosa* was evaluated by PCR, and 52.4% of the total infected plants were positive. Positive plants were then selected for bacterial population and symptom analyses 18 months after inoculation. Among them, 71% showed colonization 5 cm aip, but not in more distal parts of the plant. This indicates that bacterial migration through the xylem vessels was restrained (Fig. 7a). In contrast, WT plants and only two transgenic clones were colonized in more distal regions, at 30 cm aip (Fig. 7a). Interestingly, though, the clones V4.6 and V5.10 were the clones with higher bacterial colonization and the only ones showing symptoms, following the same pattern observed for WT plants (Fig. 7b). Nevertheless, the symptom severity was much lower than observed in WT (Fig. 7c). Curiously, transgenic lines without long-distance colonization were symptom-free during the extend of the evaluated time course of 18 months (Fig. 7c). These results suggest that the presence of *EFR* affects bacterial colonization throughout the xylem.

**Fig. 7.**
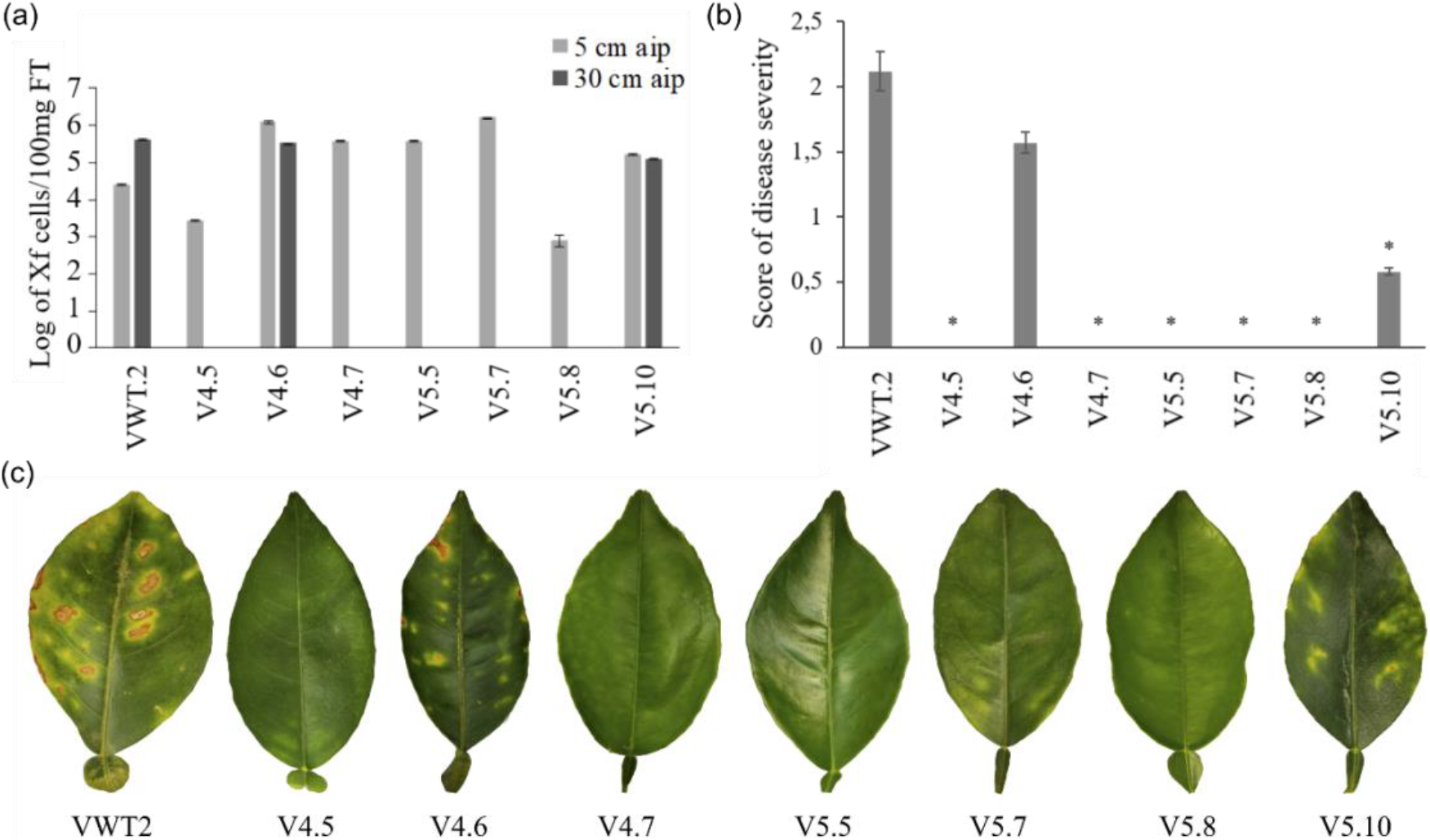
Evaluation of bacterial growth and CVC symptomatology in *EFR* transgenic sweet orange and wild-type (WT) 18 months after inoculation. **(a)** The *X. fastidiosa* population was evaluated at two different regions of the plant, 5 and 30 cm above inoculation point (aip) by TaqMan assay. **(b)** The severity of CVC leaf symptoms. **(c)** Representative leaves showing CVC symptoms. Data are expressed as the means ± standard error of three biological replicates. Statistical differences compared to WT were determined by the Student’s *t*-test (* *p* < 0.05).

## Discussion

In this study, we capitalized on observations in *Arabidopsis*, revealing EFR-dependent responses to *X. fastidiosa* and investigated whether the stable expression of *EFR* in sweet orange confers perception of EF-Tu derived from *X. citri* and *X. fastidiosa* and trigger immune responses. We demonstrated that the transgenic lines elicited conserved PTI outputs, such as ROS production, MAPK activation, and defense gene expression. Additionally, *EFR*-expressing citrus plants showed improved capacity to cope with both pathogens, culminating in reduced symptom development, and restrained pathogen colonization. This is the first report of a perennial species expressing the *EFR* receptor from *Arabidopsis* attempting to improve broad-spectrum resistance in sweet orange.

Improvements in crop fitness are increasingly desirable, especially considering the crop’s ability to overcome major biotic and abiotic stresses in an environmental-friendly fashion. Particularly for citriculture, managing CVC and citrus canker in the orchards currently depend on the control of the insect vector by regular insecticide applications, and by large amounts of copper spraying, respectively (Coletta-Filho *et al.*, 2020; Lamichhane *et al.*, 2018; Pignati *et al.*, 2017). Obtaining more resistant genotypes using biotechnology tools is an interesting strategy to anticipate plant security issues (Caserta *et al.*, 2020).

The heterologous expression of the EFR receptor, a *Brassicaceae*-specific PRR recognizing bacterial EF-Tu, is known to confer resistance and trigger multiple defense responses against several plant pathogenic bacteria. This strategy was effective in *N. benthamiana*, tomato, rice, wheat, potato, and *Medicago truncatula* (Boschi *et al.*, 2017; Kunwar *et al.*, 2018; Lacombe *et al.*, 2010; Pfeilmeier *et al.*, 2019; Schoonbeek *et al.*, 2015; Schwessinger *et al.*, 2015). The manipulation of PTI-related genetic traits can create more durable resistance and assure more sustainable productivity (Boutrot and Zipfel, 2017).

Transgenic citrus plants overexpressing the receptors FLS2 from *Arabidopsis* or Xa21 from *Oryza longistaminata* were previously shown to enhance citrus canker resistance (Hao *et al.*, 2016; Mendes *et al.*, 2010; Omar *et al.*, 2018; Shi *et al.*, 2016). In these cases, although citrus carries FLS2 orthologs, the receptor is likely to be weakly responsive or insensitive to flg22 from *X. citri* (flg_Xcc_) in sweet orange varieties (Shi *et al.*, 2015). Hamlin sweet orange and Carrizo citrange overexpressing *N. benthamiana FLS2* (*NbFLS2*) activated ROS production and defense marker genes in response to flg22_Xcc_, reducing canker susceptibility (Hao et al., 2015). Similar results were observed when Xa21 was expressed in sweet orange and mandarins (Mendes *et al.*, 2010; Omar *et al.*, 2018). These data demonstrate that sweet orange PRR-mediated signaling cascades are likely to be conserved and imply that the heterologous expression of the receptors is useful to increase basal defense responses in susceptible citrus genotypes. In contrast, although good efforts have been made to assess citrus canker immune responses, no data is available regarding the roles that PRRs play over the causal agent of CVC. The fastidious behavior of *X. fastidiosa* to produce symptoms under greenhouse conditions is demanding and requires long-term trials, making disease resistance analysis challenging.

The ability of EFR to confer broad-spectrum resistance is related to the relatively high sequence conservation of its immunogenic ligands. Agriculturally-relevant phytopathogenic bacteria such as *A. avenae*, *P. syringae*, *R. solanacearum* and *X. oryzae* showed reduced ability to cause disease after *EFR* gene transfer (Lacombe *et al.*, 2010; Lu *et al.*, 2015; Schoonbeek *et al.*, 2015; Schwessinger *et al.*, 2015). Likewise, the transfer of *EFR* to sweet orange was effective to improve *X. citri* and *X. fastidiosa* perception and thereby to enhance citrus’ immune system. The functionality of the EFR receptor was first confirmed by the ROS assay in response to elf18_Ec_ peptide (Fig. 3a). This functional conservation indicates that, even with the high evolutionary distance between *Arabidopsis* and *Citrus*, PTI cascades share co-receptors and downstream components, as previously reported for other species (Holton *et al.*, 2015; Schwessinger *et al.*, 2015). The treatment with citrus pathogen-derived elf peptides led to a slightly delayed ROS burst, and although the consistent perception of the elf18_Xcc_ occurs, mild ROS production was observed for the elf26_Xf_ peptide (Fig. 3b, c). When compared to elf18_Ec_, the peptides elf18_Xcc_ and elf18_Xf_ have 2 and 5 amino acid substitutions at the N-terminus, respectively (Fig. 1d). Although elf18_Xcc_ kept the four key conserved amino acids from the minimal peptide, the K2Q replacement in elf18_Xf_ (Fig. 1d) caused a reduction of the EC_50_ value to ~30 nM, since it is a key residue required for elf full activity (Kunze *et al.*, 2004). Here, we used elf26_Xf_, a longer version of elf peptide described to also show high reactivity with the EFR ectodomain (Kunze *et al.*, 2004). This longer acetylated version of *X. fastidiosa* EF-Tu N-terminus was more efficient in eliciting immune responses in citrus transgenic plants (Fig. 3c, f).

PRRs engage signaling components upon PAMP perception (Zipfel and Oldroyd, 2017). In agreement with established EFR-mediated signaling, seedling growth repression induced by OMVs from *X. fastidiosa* was impaired in the *bak1-5* mutant (Fig. 1b), a mutant affected in the major PRR co-receptor BAK1 (Schwessinger *et al.*, 2011). Although some co-receptors, such as BAK1, are highly conserved (91% similarity) between *Citrus* and *Arabidopsis* (Fig. S4), other PRR interactors such as BIK1 are less so, yet showing considerable sequence conservation (80% similarity) (Fig. S5). Nevertheless, functional conservation is likely to occur in different extents of BAK1 phosphorylation leading to regular MAPK activation in citrus *EFR*-transgenic plants. To verify this hypothesis, in addition to the ROS production, we evaluated well-characterized defense responses activated after PAMP recognition in citrus. The MAPK activation (Fig. 4) and the expression profile of the defense marker genes *SGT1*, *WRKY23*, *EDS1,* and *NPR2* were assessed (Shi *et al.*, 2015) (Fig. 5). MAPK phosphorylation was consistent for all elf peptides, especially after 45 minutes of peptide treatment. It is noteworthy that, when compared to V4, the V5 line showed visible stronger activation, and this transgenic event showed some extent of constitutive MAPK activation even without PAMP treatment (Fig. 4); probably as a result of ectopic EFR expression and perception of naturally occurring bacteria in the citrus phyllosphere. It has been suggested that the auto-activation of the immune system can lead to ligand-independent enhanced disease resistance (Holton *et al.*, 2015), which could be the case for the V5 line. However, in this line higher MAPK activation could be induced upon elf treatment. Noteworthy, the V5 line showed the highest increase in *X. citri* resistance, showing fewer symptoms compared to V4 and WT. When infected with *X. fastidiosa,* even when the bacteria could migrate throughout the host and produced chlorotic lesions, symptoms were milder compared to the WT plants. The differences in disease severity observed among the clones may be due to chimera, since this is a common feature in citrus transgenic plants (Caserta *et al.*, 2017; Domínguez *et al.*, 2004). This is likely to be the reason why the clones displayed variable levels of resistance to *X. fastidiosa*.

Although the induction of defense responses by *X. citri- and X. fastidiosa*-derived elf peptides was weaker than what was observed upon elf18_Ec_ treatment, when the transgenic plants were challenged with the pathogens, the impact on colonization and symptom development was evident. The *X. citri* biofilm is highly enriched in EF-Tu indicating that this protein is being recognized by EFR during bacterial growth. Noticeable citrus canker symptom reduction together with the restrained bacterial growth supports this hypothesis. Particularly, the V5 line proved to be more efficient in impairing pathogen progression, corroborating with all previous results, such as ROS production and the increased activation of MAPKs (Fig. 3e; Fig. 4).

To our knowledge, this is the first time that immune receptor mutants in the genetic model *Arabidopsis* and transgenic plants expressing a PRR receptor have been challenged with *X. fastidiosa*. This phytopathogen is a slow-growing bacterium requiring long experimental time courses of up to one year for symptom development and disease evaluation. On the other hand, *Arabidopsis* and tobacco are alternative hosts for *X. fastidiosa* and have been used to study many aspects of *Xylella*-plant host interactions (Caserta *et al.*, 2017; Ge *et al.*, 2020; Lopes *et al.*, 2020; Pereira *et al.*, 2019). Here, both *Arabidopsis* and tobacco model plants were used to determine the behavior of *X. fastidiosa* when *EFR* was knocked-out or expressed (Fig. 1, Fig. S2 and S3). The assays indicated that EFR recognizes *X. fastidiosa* and subsequently activates plant defenses since significantly less symptoms were observed in *EFR*-expressing plants (Fig. 1, Fig. S2 and S3). These results encouraged us to transform a commercial variety of sweet orange, where, in agreement with what was observed for tobacco, reduced symptom severity in transgenic lines. In addition, *X*. *fastidiosa* failed in colonizing more distal parts of the majority of *EFR* citrus transgenic lines. Even in the individual clones where bacterial colonization was higher, symptom severity was mild compared to the WT (Fig. 7b, c). It is already known that when *X. fastidiosa* reaches high populations the biofilm produced in the xylem vessels is rich in EF-Tu (Silva *et al.*, 2011). So, we hypothesized that it can be recognized by EFR and trigger PTI. The extracellular products of OMV released by *X. fastidiosa* in xylem vessels may have alternative roles that might modulate movement and biofilm formation in the host (Ionescu *et al.*, 2014). Interestingly, the OMVs produced by *X. fastidiosa* have abundant EF-Tu content (Nascimento *et al.*, 2016) likely to be involved with functions other than protein synthesis (Matsumoto *et al.*, 2012). In turn, plants can recognize immunogenic patterns present in OMVs (Bahar *et al.*, 2016), consistent with OMV-induced seedling growth repression (Fig. 1b). Therefore, we suggest that OMVs may move along the xylem vessels releasing sufficient EF-Tu to be recognized by EFR expressed in the transgenic lines. Thus, the host immune system is activated and prevents further bacterial colonization and establishment. Our results show that this approach can be an interesting strategy to improve disease resistance in agriculturally relevant species affected by *X. fastidiosa*, such as grapevine, olives, almonds, and coffee (Almeida *et al.*, 2019; Coletta-Filho *et al.*, 2020).

To the best of our knowledge, citrus lacks receptors capable of recognizing EF-Tu. Although EFR is restricted to the *Brassicaceae* family, recognition of EF-Tu by an unknown receptor has been reported in rice (Furukawa *et al.*, 2014). Although no EFR homolog was found in *Citrus* so far (Magalhães *et al.*, 2016), two LRR-RKs were highly expressed in the resistant species *C. reticulata* after *X. fastidiosa* infection (Rodrigues *et al.*, 2013). These genes were classified as belonging to the XII group of LRR-RKs (Magalhães *et al.*, 2016), and might thus be involved in the perception of some unknown *Xylella*-derived PAMP(s).

In summary, our results showed that the expression of *EFR* in sweet orange triggers ligand-dependent activation of defense responses, leading to improved resistance against major citrus bacterial pathogens. The increments on resistance aiming at avoiding or decreasing the use of agrochemical inputs is economically viable and sustainable. Although few studies have been carried out for characterization under field conditions, tomato EFR-expressing plants have been evaluated for field trials, showing promising results for bacterial disease management caused by *R. solanacearum* and *X. performans* (Kunwar *et al.*, 2018). This opens possibilities and encourages the use of PRRs to confer broad-spectrum resistance as a strategic approach that may support biotechnology citrus breeding programs.

## Acknowledgments

We thank L. F. C. Silva from the Centro de Citricultura “Sylvio Moreira” at the Instituto Agronômico de Campinas (IAC) for greenhouse assistance. This work was supported by a research grant from the Fundação de Amparo à Pesquisa do Estado de São Paulo (2013/10957-0) and INCT-Citros (Fapesp 2014/50880-0 and CNPq 465440/2014-2). LKM and DMM, Ph.D. candidates from the Graduate Program in Genetics and Molecular Biology (Unicamp), were supported by fellowships from Coordenação de Aperfeiçoamento de Pessoal de Nível Superior grant 001 and FAPESP (2013/01412-0), respectively. NSTS is a postdoctoral fellow supported by FAPESP (2019/01447-5). AAS is a recipient of research fellowships from CNPq. Work in the Zipfel laboratory on the interfamily transfer of EFR is funded by the Gatsby Charitable Foundation and the 2Blades Foundation. Research in the Robatzek laboratory is funded by the Deutsche Forschungsgemeinschaft (DFG), supporting S.R. with a Heisenberg fellowship (RO 3550/14-1) and K.R. with an accompanying grant (RO 3550/13-1).

## Author contribution statement

AAS, SR and CZ conceived and designed this research. LKM, NSTS, DMM, RRSN, and KR conducted experiments and analyzed data. LKM and NSTS drafted the manuscript. LKM, NSTS, KR, SR, AAS, and CZ contributed to the interpretation of the data and provided intellectual input. AAS and CZ provided reagents, analytical tools, and revised the manuscript. All authors read and approved the final manuscript.

## Supplementary Material

**Table S1.**
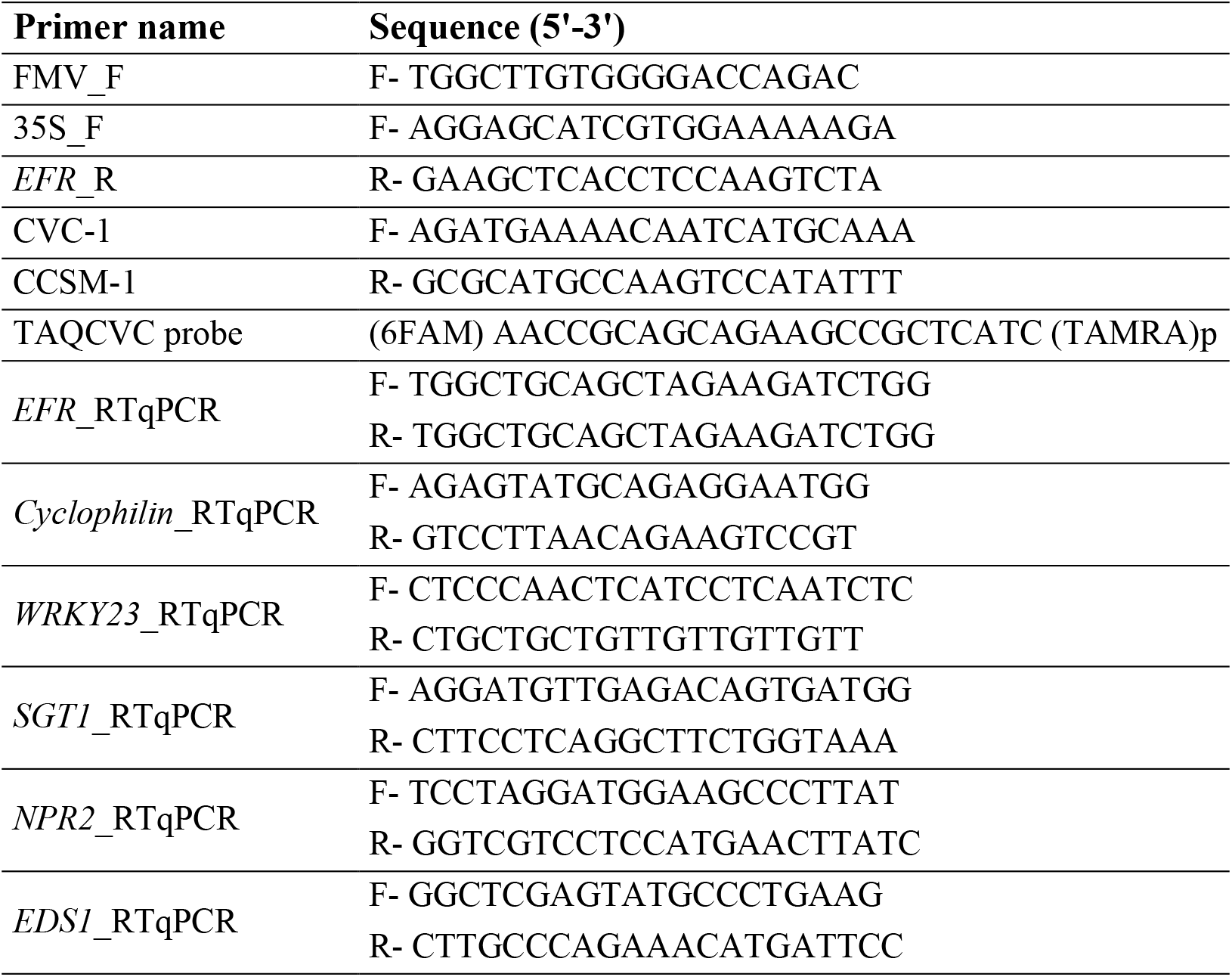
List of primers and probes used in this study.

**Table S2.** *Citrus* transformation assays performed in Valencia sweet orange for *EFR* gene transfer. Number of recovered plants and transformation efficiency are indicated.

| Experiment | N° of explants | N° of shoots | GUS (+) shoots | PCR (+) plants/<br>Analyzed plants | Transformation efficiency (%)* |
| --- | --- | --- | --- | --- | --- |
| 1 | 248 | 121 | 5 | 5/5 | 2.02 |
| 2 | 188 | 80 | 3 | 3/3 | 1.60 |
| 3 | 151 | 101 | 14 | 5/14 | 9.27 |
| 4 | 148 | 96 | 17 | 7/17 | 11.49 |

\**gus* (+) shoots/total explants infected

**Fig. S1.**
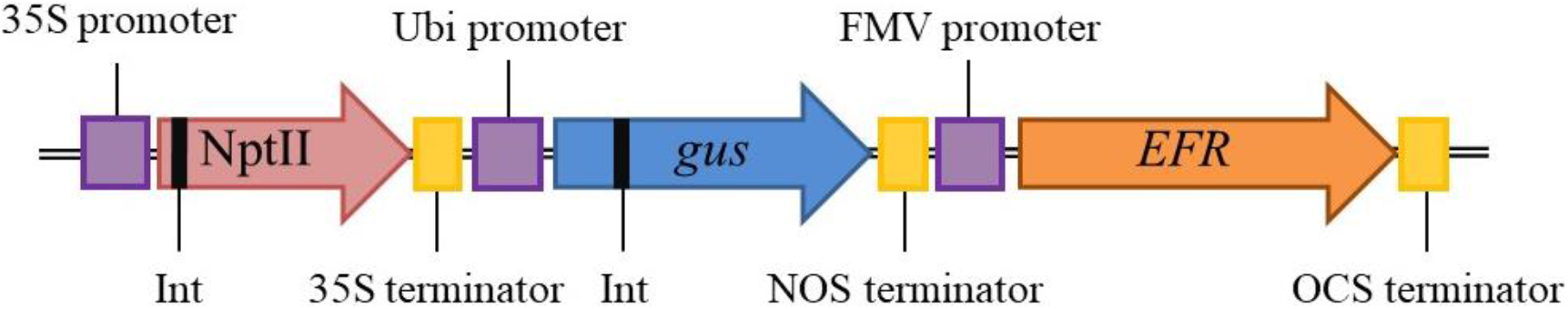
Schematic representation of the expression cassette containing the *EFR* gene from *Arabidopsis* used for sweet orange *Agrobacterium*-mediated transformation. The neomycin phosphotransferase encoding sequence (*NptII*) used as the plant selection marker is under the control of the 35S promoter and terminator. The *gus* reporter gene expression is regulated by the ubiquitin (Ubi) promoter and the nopaline synthase (NOS) terminator. The *EFR* gene is under the control of the *fig mosaic virus* (*FMV*) promoter and the *octopine synthase* (*OCS*) terminator.

**Fig. S2.**
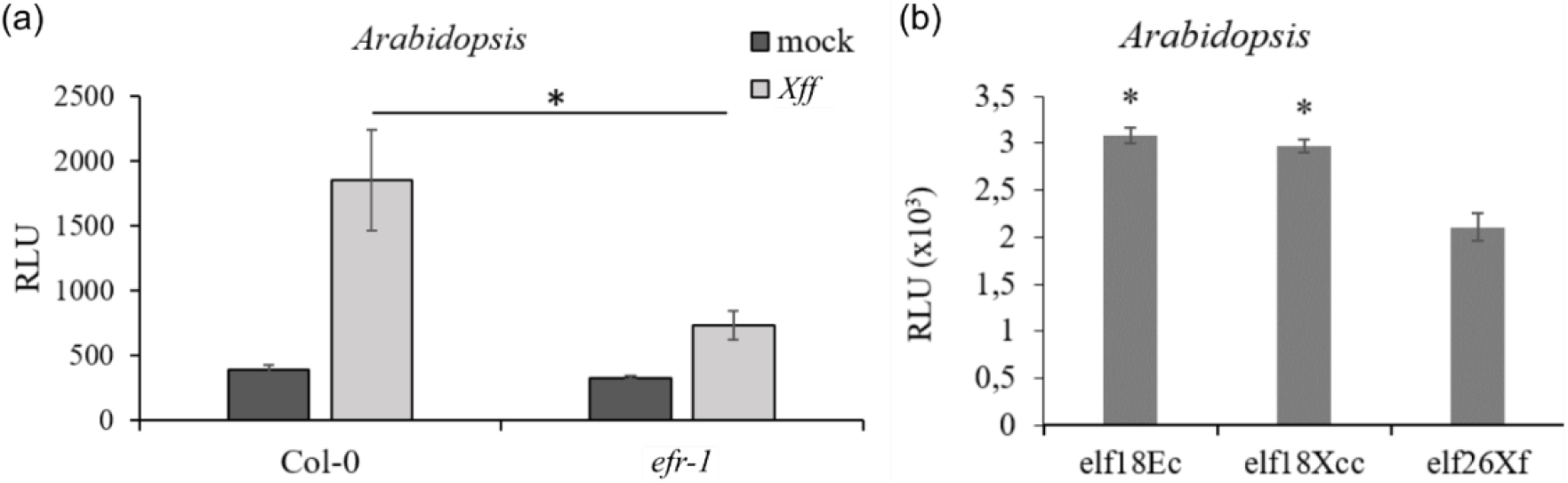
ROS burst in Col-0 and *efr-1* mutant after treatment with *Xff* (OD_600_ 0.125) and with multiple elf peptides. **(a)** Total photon count represented as relative light units following PAMP treatment (n=10) and **(b)** ROS production in *Arabidopsis* (Col-0) triggered by elf18 peptides. Error bars represent standard error of the mean. Statistical differences are represented by asterisks and were calculated using a two-tailed *t*-test (* *p* < 0.05). The experiments were performed twice with similar results.

**Fig. S3.**
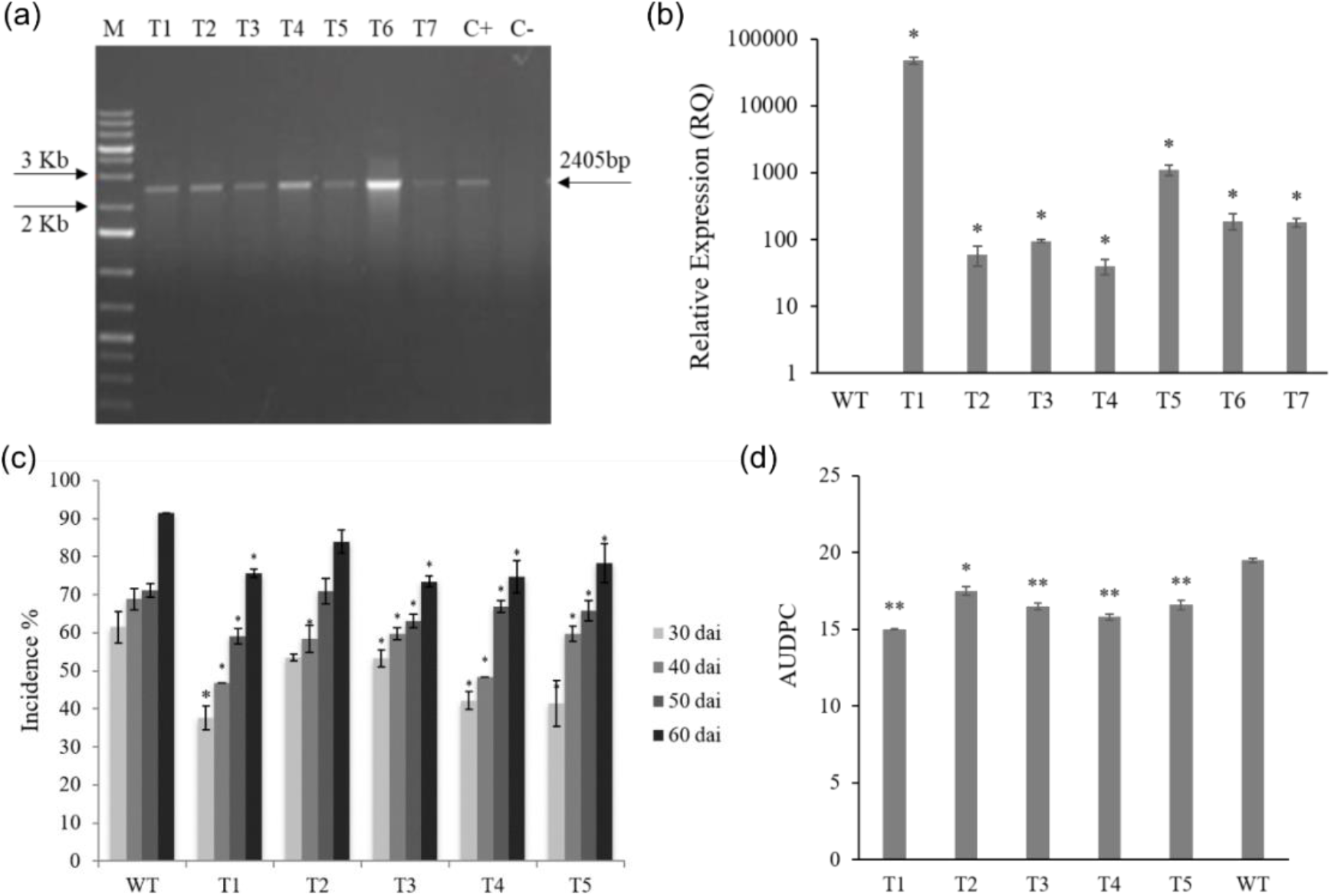
Transgenic tobacco expressing *EFR*. **(a)** PCR confirmation of five transgenic lines showing the expected amplicon of 2403 bp (black arrow). L = 1Kb DNA ladder (Thermo Fisher Scientific), T1-T5 = transgenic lines, C+ = positive control (empty vector), C- = WT tobacco genomic DNA. **(b)** Relative expression profile of the *EFR* encoding gene in transgenic tobacco plants. RLU: relative unit of light. Values are means ± standard error (SE) of at least six biological replicates. **(c)** Symptomatology of *X. fastidiosa* infection in *EFR*-expressing transgenic tobacco showing the average disease incidence in **(c)** and the area under the disease progress curve (AUDPC) in **(d)** The percentage of incidence was used to calculate the disease progression curve at 30, 40, 50 and 60 dai. Results are shown as means of at least three independent biological replicates ± standard error (SE). Statistical differences compared to WT were determined by the Student’s *t* test (* *p* < 0.05; ** *p* < 0.01).

**Fig. S4.**
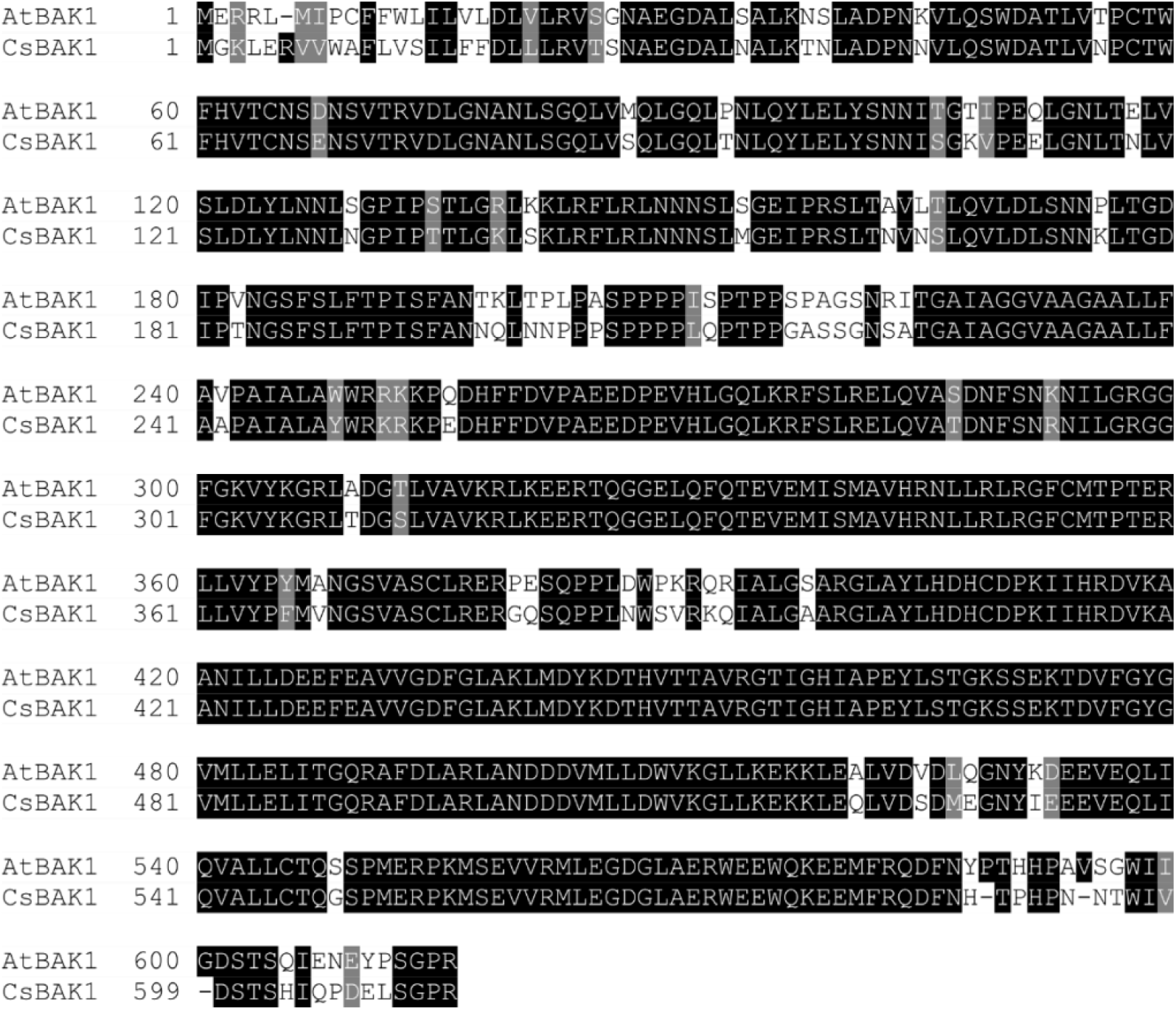
Protein sequence alignment of BRASSINOSTEROID INSENSITIVE 1-ASSOCIATED RECEPTOR KINASE 1 (BAK1) of *Arabidopsis* (AtBAK1 – NP_567920.1) and its ortholog in *C. sinensis* (CsBAK1 – XP_006493289.1). The sequences showed 99% coverage and 91% similarity. The alignment was performed at the T-Coffee server.

**Fig. S5.**
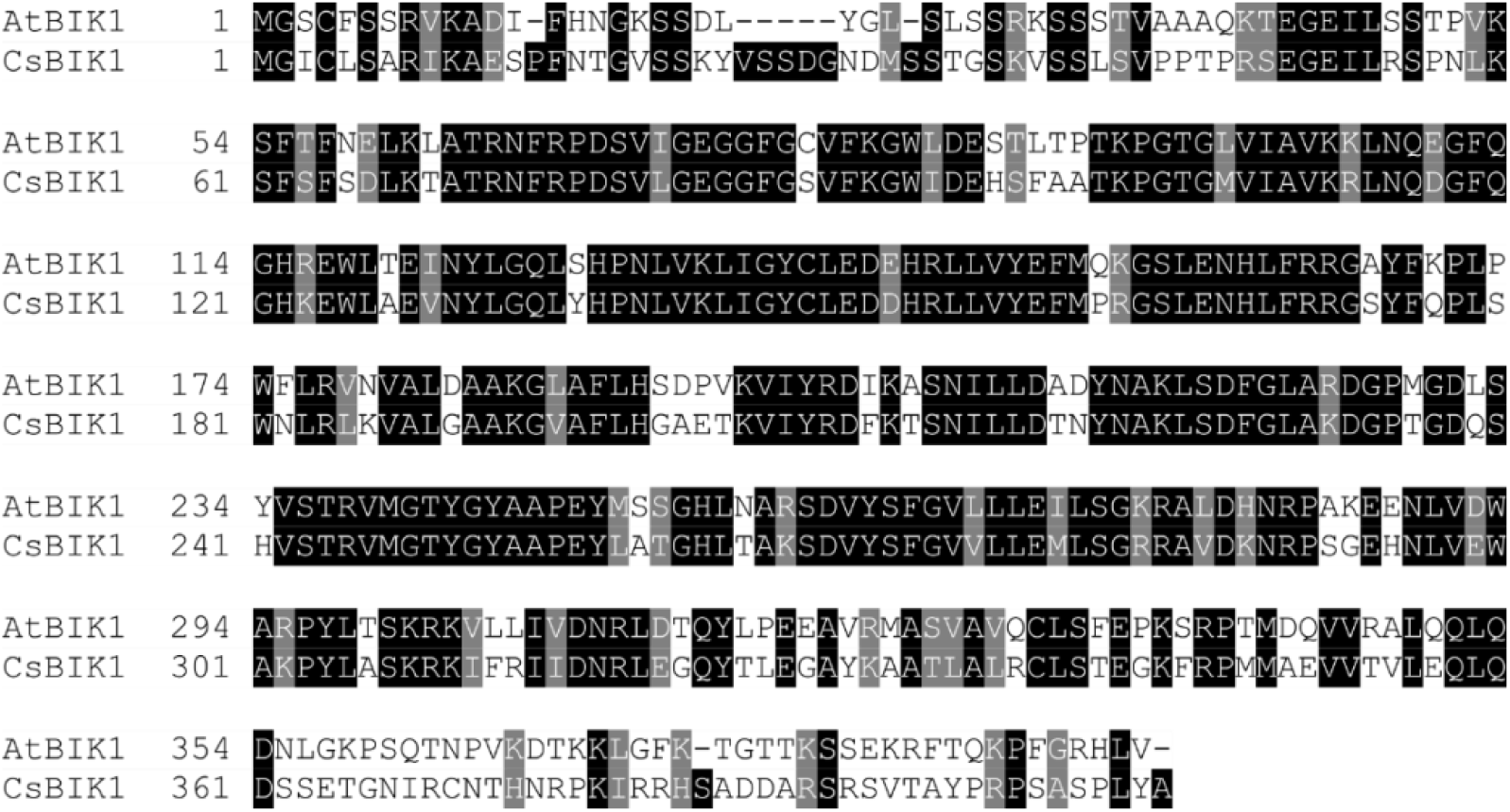
Protein sequence alignment of the serine/threonine-protein kinase BOTRYTIS-INDUCED KINASE 1 (BIK1) from *Arabidopsis* (AtBIK1 – NP_181496.1) and its ortholog from *C. sinensis* (CsBIK1 – XP_006488335.1). The sequences showed 89% coverage and 80% similarity among the amino acids. The alignment was performed at the T-Coffee server.

